# A mathematical dissection of the adaptation of cell populations to fluctuating oxygen levels

**DOI:** 10.1101/827980

**Authors:** Aleksandra Ardaševa, Robert A Gatenby, Alexander R A Anderson, Helen M Byrne, Philip K Maini, Tommaso Lorenzi

## Abstract

The disordered network of blood vessels that arises from tumour angiogenesis results in variations in the delivery of oxygen into the tumour tissue. This brings about regions of chronic hypoxia (*i.e*. sustained low oxygen levels) and regions with alternating phases of low and relatively higher oxygen levels within vascularised tumours, and makes it necessary for cancer cells to adapt to fluctuating environmental conditions. We use a phenotype-structured model to dissect the evolutionary dynamics of cell populations exposed to fluctuating oxygen levels. In this model, the phenotypic state of every cell is described by a continuous variable that provides a simple representation of its metabolic phenotype, ranging from fully oxidative to fully glycolytic, and cells are grouped into two competing populations that undergo heritable, spontaneous phenotypic variations at different rates. Model simulations indicate that, depending on the rate at which oxygen is consumed by the cells, nonlinear dynamic interactions between cells and oxygen can stimulate chronic hypoxia and cycling hypoxia. Moreover, the model supports the idea that under chronic-hypoxic conditions lower rates of phenotypic variation lead to a competitive advantage, whereas higher rates of phenotypic variation can confer a competitive advantage under cycling-hypoxic conditions. In the latter case, the numerical results obtained show that bet-hedging evolutionary strategies, whereby cells switch between oxidative and glycolytic phenotypes, can spontaneously emerge. We explain how these results can shed light on the evolutionary process that may underpin the emergence of phenotypic heterogeneity in vascularised tumours.

## 1 Introduction

Marked spatial variations in the molecular properties of clinical cancers have been well recognised. This is often ascribed to evolution driven by genetic mutations (‘branching clonal evolution’). An alternative hypothesis is that the cancer cells are simply evolving to adapt to spatial and temporal variations in microenvironmental conditions that result from heterogeneous blood flow. The vascular structure in tumours is highly disordered and is constantly re-modelled via processes such as sprouting angiogenesis (*i.e*. formation of new blood vessels from pre-existing ones) and vascular regression and dilation (Carmeliet and Jain, 2000; Welter and Rieger, 2012). As a consequence, there can be spatial and temporal variations in the delivery of oxygen to tumour regions, leading to oscillations between phases of oxygen-deprivation and re-oxygenation (Kimura et al., 1996).

Experimental and clinical studies have shown that these oscillations can occur on a variety of timescales, ranging from minutes to weeks (Carmeliet and Jain, 2000; Dewhirst, 2009), and may lead to the emergence of regions of normoxia (*i.e*. high oxygen levels), chronic hypoxia (*i.e*. sustained low oxygen levels) and regions with fluctuating levels of oxygen (*i.e*. transient periods of low and relatively higher oxygen levels) within vascularised tumours (Matsumoto et al., 2010; Michiels et al., 2016; Ron et al., 2019).

The evolutionary consequences of these spatial and temporal variations can be profound. Clearly, optimal fitness for cancer cells in a poorly perfused region that is hypoxic, acidic and lacks growth factors requires a different phenotype compared to a cancer cell in a well-perfused physiological environment. Furthermore, rapid, stochastic changes in environmental conditions apply additional selection forces. Here, cells must be capable of rapidly adapting to unpredictable and potentially lethal environmental conditions. Furthermore, hypoxic and acidic environments generate genotoxic environments and the transition from hypoxic to normoxic conditions can generate bursts of oxygen free radicals that can induce widespread tissue and cellular damage.

Previous empirical and theoretical work has suggested that temporal variations in oxygen levels make it necessary for cancer cells to adapt to fluctuating environmental conditions (Gillies et al., 2018; Amend et al., 2018) can substantially impact evolutionary dynamics of cancer populations by increasing clonal diversity, promoting metastasis and supporting more plastic phenotypic variants (Cairns et al., 2001; Cairns and Hill, 2004; Louie et al., 2010; Verduzco et al., 2015; Chen et al., 2018; Saxena and Jolly, 2019). In particular, it has been hypothesised that – by analogy with bacterial populations facing unpredictable environmental changes (Kussell and Leibler, 2005; Smits et al., 2006; Veening et al., 2008; Acar et al., 2008; Beaumont et al., 2009; Nichol et al., 2016) – cancer cell populations could utilise risk spreading through stochastic phenotype switching, which is also known as bet-hedging (Philippi and Seger, 1989), as an adaptive strategy to survive in the harsh, constantly changing environmental conditions associated with intermittent hypoxia (Gravenmier et al., 2018; Gillies et al., 2018).

In this paper, we use a phenotype-structured model of evolutionary dynamics in time-varying but spatially homogeneous environments to elucidate the mechanisms that underpin the adaptation of cell populations to fluctuating oxygen inflow. Building upon the modelling framework that we presented in Ardaševa et al. (2020), the model is defined in terms of a system of non-local parabolic partial differential equations (PDEs) for the evolution of the phenotype distributions of two competing cell populations that undergo heritable, spontaneous phenotypic variations at different rates. Similar PDEs modelling the evolutionary dynamics of populations structured by continuous traits in periodically-fluctuating environments have recently received increasing attention from the mathematical community (Lorenzi et al., 2015; Mirrahimi et al., 2015; Iglesias and Mirrahimi, 2018; Carrere and Nadin, 2019).

In the model considered here, the phenotypic state of every cell is modelled by a continuous variable that provides a simple representation of its metabolic phenotype, ranging from oxidative to glycolytic. The phenotypic fitness landscape of the two cell populations evolves in time due to variations in the concentration of oxygen. The oxygen concentration is governed by an ordinary differential equation (ODE) with a time-dependent source term that models the effect of variations in the oxygen supply. The fact that oxygen is consumed by the cells is taken into account by a negative term coupling the ODE with the system of PDEs.

The paper is organised as follows. In Section 2, we introduce the equations of the model and the underlying modelling assumptions. In Section 3, we present the main numerical results of our study complemented by analytical results obtained for a model corresponding to a simplified scenario, and discuss their biological relevance. In Section 4, we explain how these mathematical results can shed light on the evolutionary process that underpins the emergence of phenotypic heterogeneity in vascularised tumours. Section 5 concludes the paper and provides a brief overview of possible research perspectives.

## 2 Description of the model

We study the evolutionary dynamics of two competing cell populations in a well-mixed system. Cells proliferate (*i.e*. divide and die) and undergo spontaneous, heritable phenotypic variations. We assume the two populations differ only in their rate of phenotypic variation. The population undergoing phenotypic variations at a higher rate is labelled by the letter *H*, while the other population is labelled by the letter *L*.

As summarised by the schematic in Fig. 1A, we represent the phenotypic state of every cell by a continuous variable *x* ∈ [0, 1]. In particular, we assume that: cells in the phenotypic state *x* = 0 have a fully oxidative metabolism and produce energy through aerobic respiration only; cells in the phenotypic state *x* = 1 express a fully glycolytic metabolism and produce energy through anaerobic glycolysis only; cells in other phenotypic states *x* ∈ (0, 1) produce energy via aerobic respiration and anaerobic glycolysis, and higher values of *x* correlate with a less oxidative and more glycolytic metabolism.

**Fig. 1.**
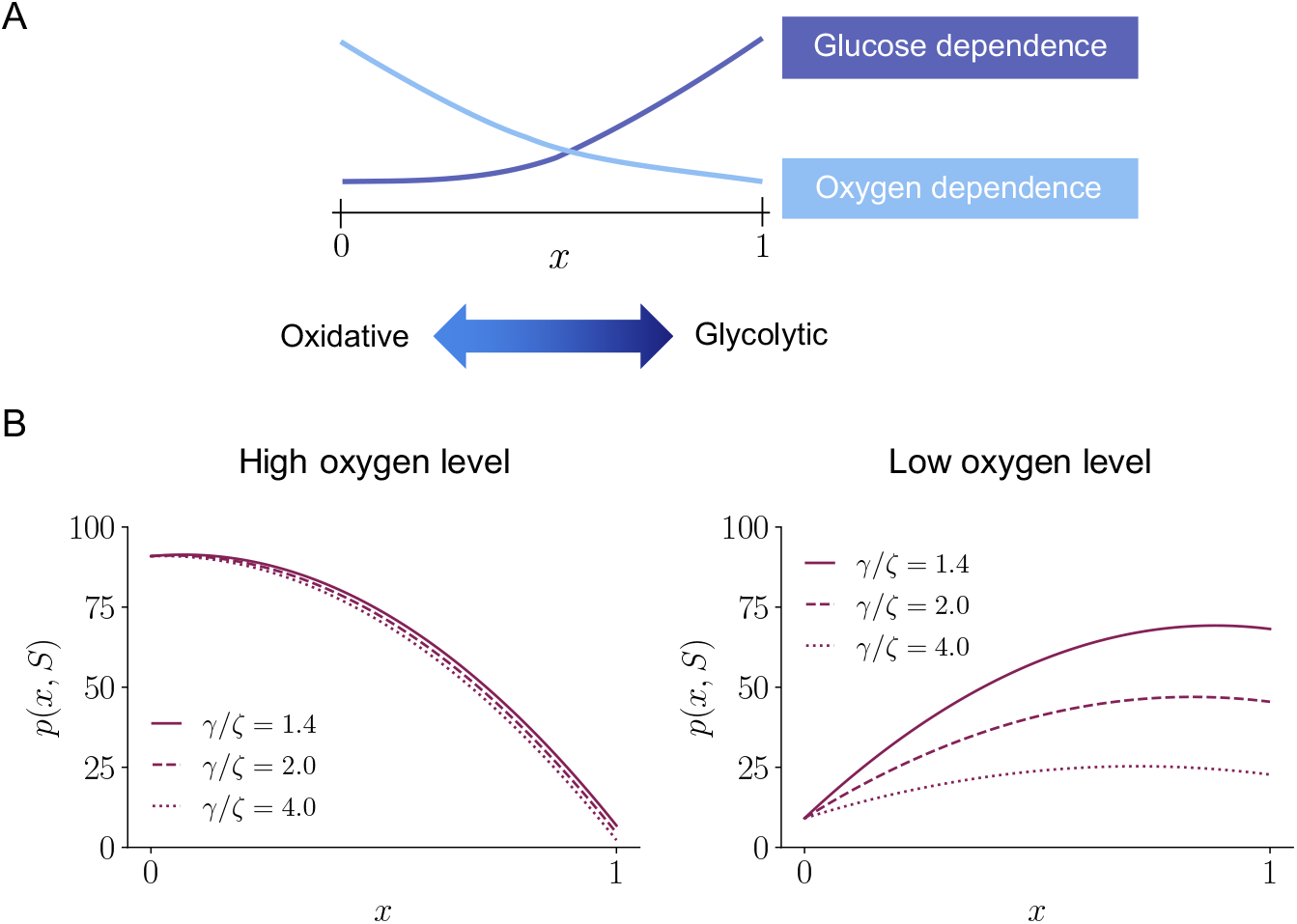
**A**. Schematic diagram illustrating the relationship between the variable *x* ∈ [0, 1], the cell phenotype and the dependence on oxygen and glucose of energy production for different phenotypic variants. **B**. Plot of the cell division rate *p*(*x, S*) defined according to Eqn. (7) in the case of a relatively high oxygen level (*i.e. S* = 10) and a relatively low oxygen level (*i.e. S* = 0.1), for increasing values of the fitness cost associated with glycolytic metabolism (*i.e*. increasing values of the quotient *γ/ζ* ≥ 1).

The oxygen concentration in the system at time *t* ∈ [0, ∞) is denoted by *S*(*t*). Based on the observation that glucose levels in biological tissues are usually high enough not to represent a limiting factor for the proliferation of cells (Gravenmier et al., 2018), for the sake of simplicity, we do not model the dynamics of the glucose concentration.

We describe the phenotype distributions of the two cell populations at time *t* by means of the population density functions *n_H_*(*x, t*) and *n_L_*(*x, t*). We define the size of populations *H* and *L*, and the total number of cells inside the system at time *t*, respectively, as

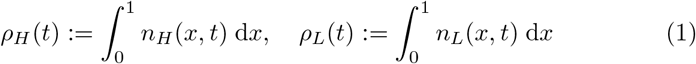

and

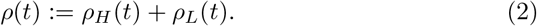

Moreover, we define the mean phenotype and the phenotypic variance of population *i* ∈ {*H, L*} at time *t*, respectively, as

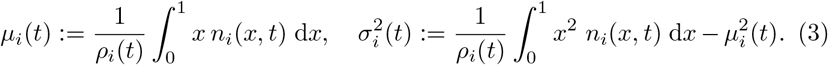

### 2.1 Cell dynamics

Building upon the modelling framework that we presented in Ardaševa et al. (2020), we describe the evolution of the two cell populations through the following system of conservation equations for the population density functions

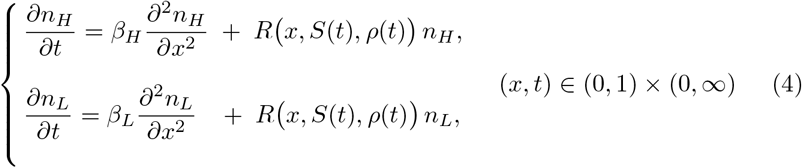

subject to no-flux boundary conditions, *i.e*.

In the non-local parabolic PDEs (4), the diffusion terms model the effect of heritable, spontaneous phenotypic variations, which occur at rates *β_H_* and *β_L_* with

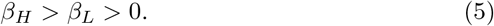

The function *R*(*x, S*(*t*), *ρ*(*t*)) represents the fitness of cells in the phenotypic state *x* at time *t* under the environmental conditions given by the oxygen concentration *S*(*t*) and the total number of cells *ρ*(*t*). This function can be seen as the phenotypic fitness landscape of the two cell populations at time *t*. We use the following definition

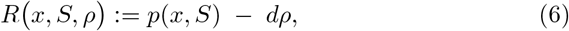

where *p*(*x, S*) is the division rate of cells in the phenotypic state *x* under the oxygen concentration *S*, while the term *dρ*, with *d* > 0, models the rate of cell death due to intrapopulation and interpopulation competition. In order to model the fact that fully oxidative phenotypic variants (*i.e*. cells in the phenotypic state *x* = 0) have the highest fitness if oxygen is abundant (*i.e*. when *S* → ∞), whereas fully glycolytic phenotypic variants (*i.e*. cells in the phenotypic state *x* = 1) are the fittest in hypoxic conditions (*i.e*. when *S* → 0), we define the cell division rate as

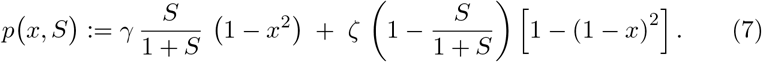

Here, the parameters *γ* and *ζ* are the maximum cell division rates of fully oxidative and fully glycolytic phenotypic variants, respectively. As we noted in Ardaševa et al. (2020), definition (7) leads to a fitness function that is close to the approximate fitness landscapes which can be inferred from experimental data through regression techniques – see, for instance, equation (1) in Otwinowski and Plotkin (2014). In fact, definition (7) can be rewritten aswhere

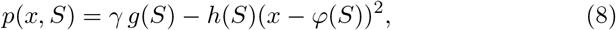

where

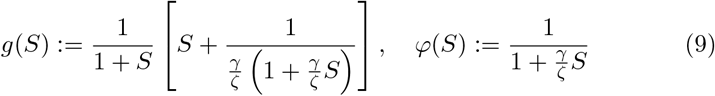

and

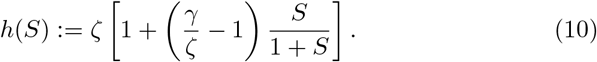

Since

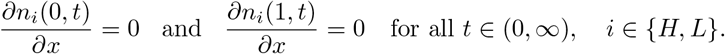

*γg*(*S*) is the maximum fitness, *φ*(*S*) is the fittest phenotypic state and *h*(*S*) is a nonlinear selection gradient. Notice that, consistent with our modelling assumptions, *φ*: [0, ∞) → [0, 1] and *φ′* < 0, so that

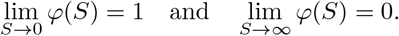

To incorporate into the model the fitness cost associated with a less efficient glycolytic metabolism (Basanta et al., 2008), we assume that

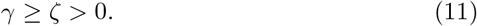

A sample of plots of the function *p*(*x, S*) for different values of the oxygen concentration *S* and of the quotient *γ/ζ* is displayed in Fig. 1B. If *γ/ζ* = 1 then there is no fitness cost associated with glycolytic metabolism, whereas increasing values of *γ/ζ* > 1 correspond to larger fitness costs of glycolytic metabolism.

### 2.2 Oxygen dynamics

We describe the oxygen dynamics via the following conservation equation for *S*(*t*):

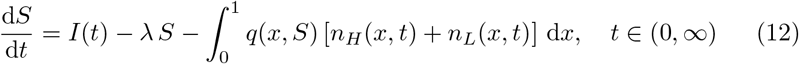

which is coupled with the non-local PDEs (4). In the ODE (12), the parameter λ > 0 represents the rate of natural decay of oxygen and the non-negative function *I*(*t*) models the rate at which oxygen is supplied to the system. The last term on the right-hand side of (12) models the rate of oxygen consumption by the cells. Here, the non-negative function *q*(*x, S*) is the consumption rate of cells in phenotypic state *x*, and we take it to be

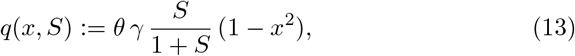

based on the following argument. Cells in the phenotypic state *x* = 1 (*i.e*. fully glycolytic phenotype) produce energy through anaerobic glycolysis only and, therefore, they do not consume any oxygen (*i.e. q*(1, *S*) = 0 for any *S*). Moreover, cells in the phenotypic state *x* = 0 (*i.e*. fully oxidative phenotype) consume oxygen at a rate proportional to their division rate, with constant of proportionality *θ* > 0 (*i.e*. 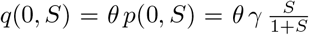). Finally, the rate at which oxygen is consumed by cells in phenotypic states *x* ∈ (0, 1) is a fraction of the consumption rate of cells in the phenotypic state *x* = 0, and higher values of *x* correlate with lower oxygen consumption (*i.e. q*(*x, S*) = *q*(0, *S*)(1 − *x*^2^) for *x* ∈ (0, 1)).

## 3 Main results

In this section, we present the results of numerical simulations of the mathematical model defined by the non-local PDEs (4) coupled with the ODE (12). We combine these numerical results with analytical results obtained for a simplified version of the model presented in Appendix A, and we discuss their biological relevance. In more detail, Section 3.1 provides a description of the numerical methods employed and the set-up of numerical simulations. In Section 3.2, we consider the case where the inflow of oxygen is constant, *i.e*. we assume

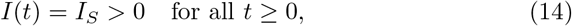

while in Section 3.3 we study the case where the oxygen inflow undergoes periodic oscillations of period *T* > 0, *i.e*. we assume

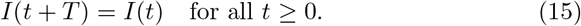

In particular, to construct numerical solutions we consider the case where

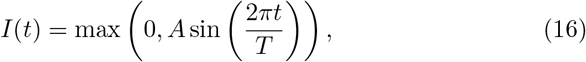

with *A* > 0 modelling the amplitude of the periodic fluctuations in oxygen inflow. Definition (16) corresponds to a biological scenario in which oxygen inflow is periodically interrupted due to, for instance, the periodic blockage of a blood vessel.

### 3.1 Numerical methods and set-up of numerical simulations

We use a uniform discretisation consisting of 200 points on the interval [0, 1] as the computational domain of the independent variable *x*. We assume *t* ∈ [0, *t_f_*], with *t_f_* = 40 being the final time of simulations, and we discretise the time interval [0, *t_f_*] with the uniform step *Δt* = 0.0001. The method for constructing numerical solutions to the system of non-local parabolic PDEs (4) subject to no-flux boundary conditions is based on a three-point finite difference explicit scheme for the diffusion terms and an explicit finite difference scheme for the reaction terms (LeVeque, 2007). Moreover, numerical solutions to the ODE (12) are constructed using the explicit Euler method.

We assume the nutrient concentration to be non-dimensionalised and use the dimensionless parameter values listed in Table 1 to carry out numerical simulations. In summary, we define the rates of phenotypic variation *β_H_* and *β_L_* so that they are consistent with typical times required by cells to acquire a glycolytic phenotype through epigenetic changes (Baumann et al., 2007). Moreover, we choose the value of the maximum cell division rate of fully oxidative phenotypic variants *γ* such that *γ* ≫ *β_H_*, in order to capture the fact that phenotypic variations occur on a slower time scale than cell division. Furthermore, to explore the effect of the cost of glycolytic metabolism on the evolutionary dynamics of the cells and on the dynamics of oxygen, we consider different values of *ζ* such that *γ/ζ* ∈ [1, 4]. Given the values of the parameters *γ*, *β_H_* and *β_L_*, we fix the value of the death rate due to competition, *d*, to be such that the long-term limit of the size of population in presence of a constant and relatively high supply of oxygen is approximatively 10^4^, which is consistent with biological data on *in vitro* cell populations (Voorde et al., 2019). Since the rate at which cells consume oxygen varies between cell lines and depends on a variety of environmental factors, including the pH level (Casciari et al., 1992), we consider a range of values for the rate of consumption of oxygen, *θ*, that is, *θ* ∈ [10^-5^, 10^-3^], to investigate also the influence this parameter has on the cell and oxygen dynamics. Finally, we choose the value of the rate of natural decay of oxygen, λ, to be consistent with values used by other authors, such as Macklin et al. (2009).

**Table 1.** Parameter values used in numerical simulations.

|  | Description | Value range |
| --- | --- | --- |
| $\beta_H$ | Rate of phenotypic variation of cells in population $H$ | $2.5 \times 10^{-2}$ |
| $\beta_L$ | Rate of phenotypic variation of cells in population $L$ | $10^{-2}$ |
| $\gamma$ | Maximum cell division rate of fully oxidative phenotypic variants | 100 |
| $\zeta$ | Maximum cell division rate of fully glycolytic phenotypic variants | [25,100] |
| $d$ | Death rate due to competition | $10^{-2}$ |
| $\theta$ | Consumption rate of oxygen | $[10^{-5}, 10^{-3}]$ |
| $\lambda$ | Rate of natural decay of oxygen | $10^{-4}$ |

We let the initial cell population density functions *n_i_*(*x*, 0) with *i* ∈ {*H, L*} be Gaussian-like functions such that

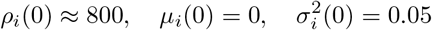

and define the initial oxygen concentration as *S*(0) = *I*(0).

### 3.2 Constant oxygen inflow

The numerical solutions presented in Fig. 2 show that when the oxygen inflow is constant [*i.e*. when the function *I*(*t*) is defined according to (14)], cell population *L* outcompetes cell population *H*, which eventually goes extinct. Moreover, the population density function *n_L_*(*x, t*) is unimodal, attaining its maximum at the mean phenotype. Further, since the oxygen concentration *S*(*t*) converges to an equilibrium value, the population size *ρ_L_*(*t*) also converges to an equilibrium value. The equilibrium value of *ρ_L_*(*t*) is approximately equal to the asymptotic value 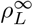 given by definition (A.6) in Appendix A, which is obtained by studying the long-time behaviour of the solutions to a simplified version of the model (*vid*. Theorem 1 in Appendix A). This is consistent with the analytical results that we presented in Ardaševa et al. (2020).

**Fig. 2.**
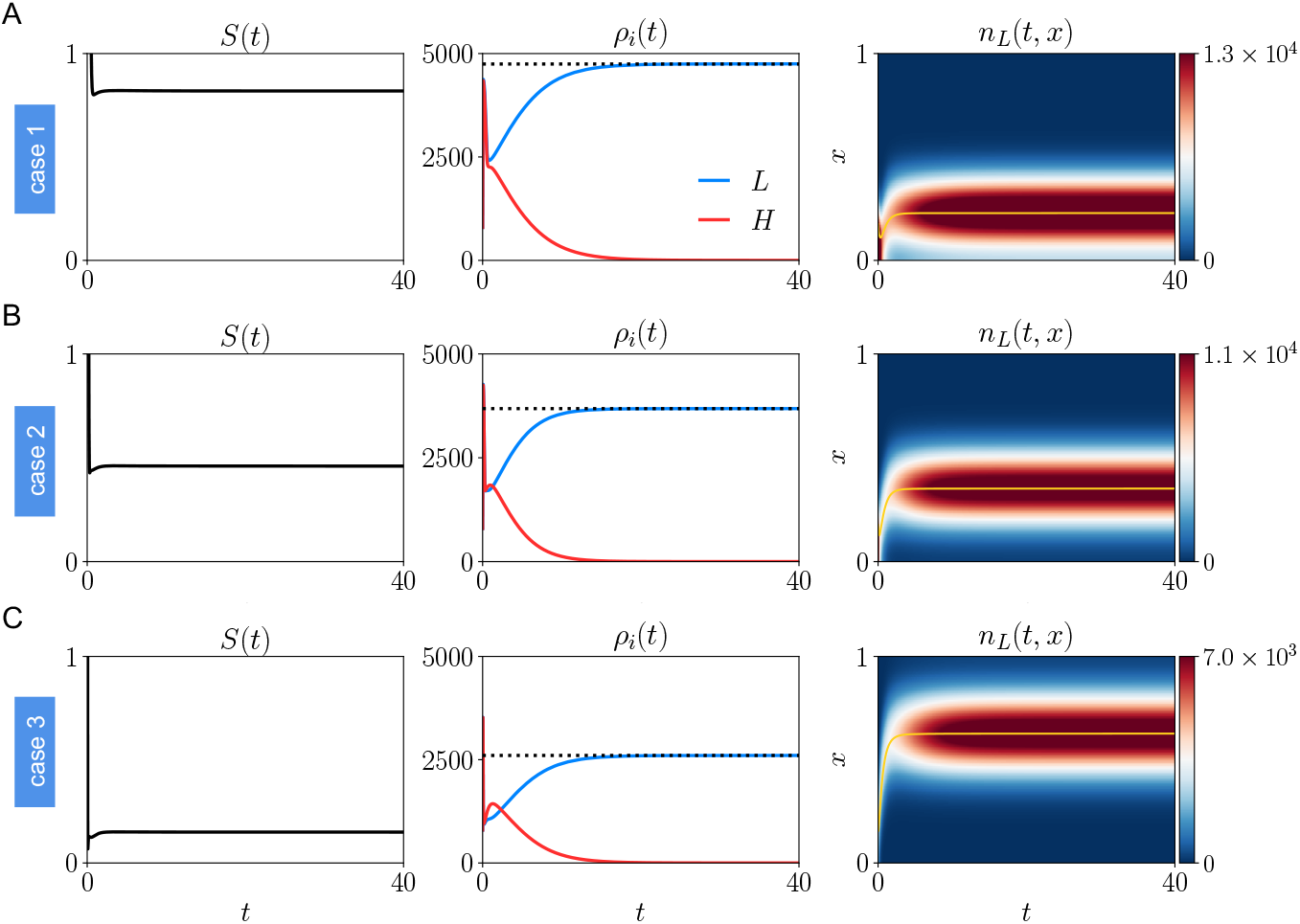
**A**. Dynamics of the oxygen concentration *S*(*t*) (first column), the population sizes *ρ_H_*(*t*) (second column, red line) and *ρ_L_*(*t*) (second column, blue line), and the population density function *n_L_*(*t, x*) (third column) obtained by solving numerically Eqns. (4) and (12), for the oxygen inflow *I*(*t*) defined via Eqn. (14) with *I_S_* = 10. The dotted lines in the second column highlight the asymptotic value 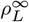 given by definition (A.6) in Appendix A, while the yellow lines in the third column highlight the mean phenotype *μ_L_*(*t*). The consumption rate of oxygen is *θ* = 5 × 10^-5^, the maximum cell division rate of fully glycolytic phenotypic variants is *ζ* = 25, and the values of the other parameters are defined as in Table 1. **B, C**. Same as row A but for *θ* = 10^-4^ (row **B**) and *θ* = 5 × 10^-4^ (row **C**).

The results displayed in Fig. 2 also show that larger values of the oxygen consumption rate *θ* lead to smaller equilibrium values of the oxygen concentration S and, therefore, smaller final values of *ρ_L_* and larger final values of *μ_L_*. Moreover, the numerical results summarised by the plots in Fig. 3 demonstrate that larger values of the fitness cost associated with glycolytic metabolism, *γ/ζ*, correspond to smaller final values of *ρ_L_* and *μ_L_*. The plots in Fig. 3 also show that lower values of *I_S_*, which lead to smaller equilibrium values of *S* for a given value of *θ* (data not shown), correlate with a weaker impact of the value of the quotient *γ/ζ* on the final values of *ρ_L_* and *μ_L_*. All these findings are consistent with the way in which the equilibrium values of the population size, 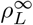, and the mean phenotype, 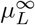, obtained through the analysis of the simplified model considered in Appendix A depend on the equilibrium value of the oxygen concentration, *S*^∞^, and on the quotient *γ/ζ* (*vid*. Theorem 1 in Appendix A).

**Fig. 3.**
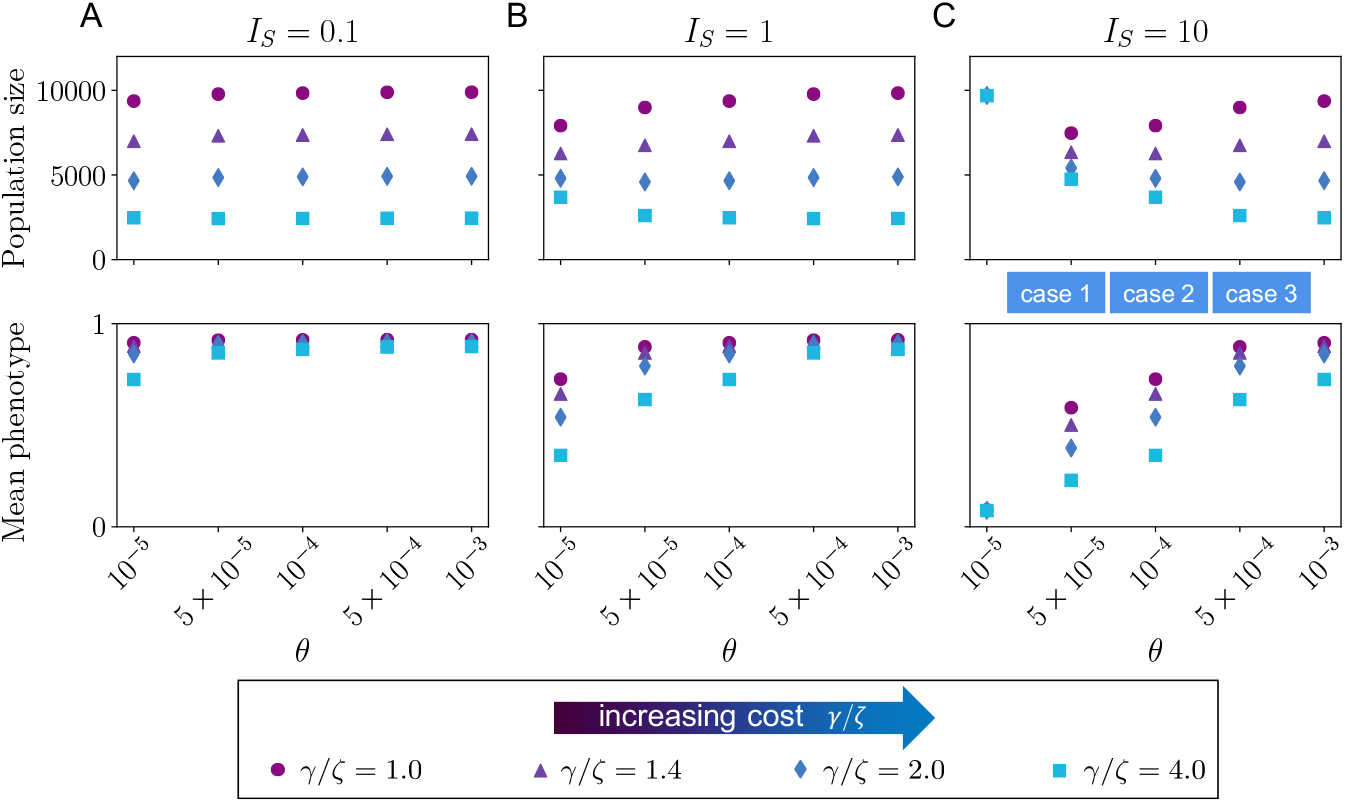
**A**. Values of the population size *p_L_*(*t*) and the mean phenotype *μ_L_*(*t*) at *t* = 40 (*i.e*. at the end of numerical simulations) obtained by solving numerically Eqns. (4) and (12), for the oxygen inflow *I*(*t*) defined via Eqn. (14) with *I_S_* = 0.1 and for different, values of the consumption rate of oxygen, *θ*, and different, values of the cost, associated with glycolytic metabolism, *γ/ζ*, obtained by changing the value of the maximum cell division rate of fully glycolytic phenotypic variants, *ζ*, and keeping the value the maximum cell division rate of fully oxidative phenotypic variants, *γ*, constant, (*i.e. γ* = 100). The values of the other parameters are defined as in Table 1. **B, C**. Same as column **A** but. for *I_S_* = 1 (column **B**) and *I_S_* = 10 (column **C**). The blue boxes in the last, panel highlight, the values of *θ* corresponding to Case 1,2,3 in Fig. 2.

Taken together, these results indicate that lower rates of heritable, sponteneous phenotypic variation constitute a source of competitive advantage under constant oxygen inflow. Furthermore, the negative feedback that regulates the growth of cell populations through oxygen consumption shapes, in a nonlinear way, the evolutionary dynamics of the cells. In particular, larger values of the rate of oxygen consumption, *θ*, lead to the emergence of lower oxygenated environments whereby phenotypic variants that rely to a larger extent on anaerobic glycolysis for energy production are ultimately selected. Finally, all other things being equal, larger values of the fitness cost associated with glycolytic metabolism, *γ/ζ*, are to be expected to promote the selection of less glycolytic phenotypic variants and to reduce the equilibrium size of cell populations exposed to constant oxygen inflow.

### 3.3 Periodic oxygen inflow

The numerical solutions presented in Fig. 4 show how the system evolves when the oxygen inflow undergoes periodic oscillations [*e.g*. when the function *I*(*t*) is defined according to (16)]: if the oxygen concentration is relatively stable (low-amplitude oscillations), cell population *L* outcompetes cell population *H*, which eventually goes extinct; if the oxygen concentration undergoes drastic, high-amplitude variations, then cell population *L* is outcompeted by cell population *H* and ultimately goes extinct. Moreover, the population density function of the surviving cell population, *n_i_*(*x, t*), is unimoda.l with maximum at the mean phenotype. Since the oxygen concentration *S*(*t*) becomes T-periodic, after an initial transient, the population size *ρ_i_*(*t*) of the surviving population also converges to a. T-periodic function. Such a. T-periodic function is approximately equal to the solution 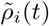 of the problem (A.11) in Appendix A, which is obtained by studying the long-time behaviour of the solutions to a. simplified version of the model (*vid*. Theorem 2 in Appendix A). This is in line with the analytical results presented in Ardaševa. et a.i. (2020).

**Fig. 4.**
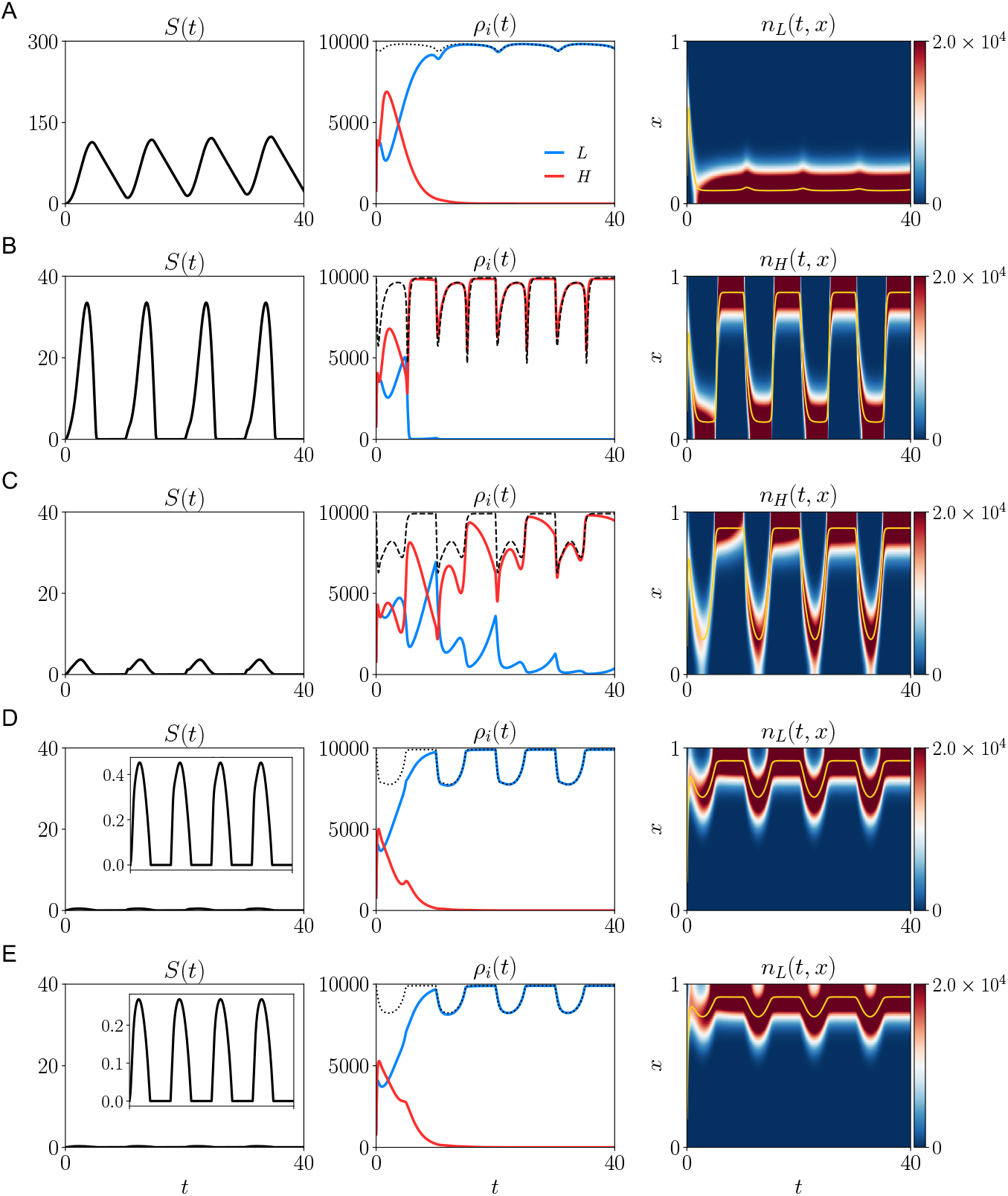
**A**. Dynamics of the oxygen concentration *S*(*t*) (first column), the population sizes *ρ_H_*(*t*) (second column, red line) and *ρ_L_*(*t*) (second column, blue line), and the population density function of the surviving population *n_i_*(*t, x*) (third column) obtained by solving numerically Eqns. (4) and (12), for the oxygen inflow *I*(*t*) defined via Eqn. (16) with *A* = 60 and *T* = 10. The dotted (or dashed) lines in the second column highlight the *T*-periodic solution 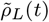 (or 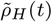) of the problem (A.11) in Appendix A, while the yellow lines in the third column highlight the mean phenotype of the surviving population *μ_i_*(*t*). The consumption rate of oxygen is *θ* = 2 × 10^-5^, the maximum cell division rate of fully glycolytic phenotypic variants is *ζ* = 7, and the values of the other parameters are defined as in Table 1. **B - E**. Same as row **A** but for *θ* = 5 × 10^-5^ (row **B**), *θ* = 10^-4^ (row **C**), *θ* = 5 × 10^-4^ (row **D**) and *θ* = 10^-3^ (row **E**).

The results displayed in Fig. 4 also show that the consumption rate of oxygen, *θ*, has a crucial impact on the dynamics of the oxygen concentration *S*(*t*) and, therefore, on the outcome of the competition between the two cell populations. In fact, *ceteris paribus*, for sufficiently small (*cf*. Fig. 4A) or sufficiently large (*cf*. Figs. 4D and 4E) values of *θ* the function *S*(*t*) is bounded well above zero or undergoes small oscillations while remaining close to zero, respectively. This brings about relatively stable oxygen concentrations in presence of which cell population *L* outcompetes cell population *H*. On the other hand, for intermediate values of *θ* (*cf*. Figs. 4B and 4C) the function *S*(*t*) oscillates between small and relatively larger values. This results in more drastic variations of the oxygen concentration, which lead cell population *L* being outcompeted by cell population *H*. As we would expect, when *S*(*t*) remains away from zero or undergoes small oscillations while remaining close to zero, the mean phenotype of the surviving population *μ_L_*(*t*) undergoes small oscillations and its value remains close, respectively, either to the fully oxidative phenotypic state *x* =0 (*cf*. Fig. 4A) or to the fully glycolytic phenotypic state *x* = 1 (*cf*. Figs. 4D and 4E). By contrast, when *S*(*t*) oscillates between small and relatively larger values, the mean phenotype of the surviving population *μ_H_*(*t*) undergoes rapid and large amplitude transitions between phenotypic states closer to *x* = 0 and phenotypic states closer to *x* = 1 (*cf*. Figs. 4B and 4C).

The numerical results in Fig. 4 refer to the case where there is no cost associated with glycolytic metabolism (*i.e. γ/ζ* = 1) and both the amplitude *A* and the period *T* of the fluctuations in oxygen inflow in definition (16) are relatively large. However, the numerical results summarised by the plots in Fig. 5 demonstrate that similar conclusions about how the oxygen consumption rate *θ* affects the outcome of the competition between the two cell populations hold when different values of the parameters *γ/ζ*, *A* and *T* are considered, provided that the value of *A* is sufficiently large.

**Fig. 5.**
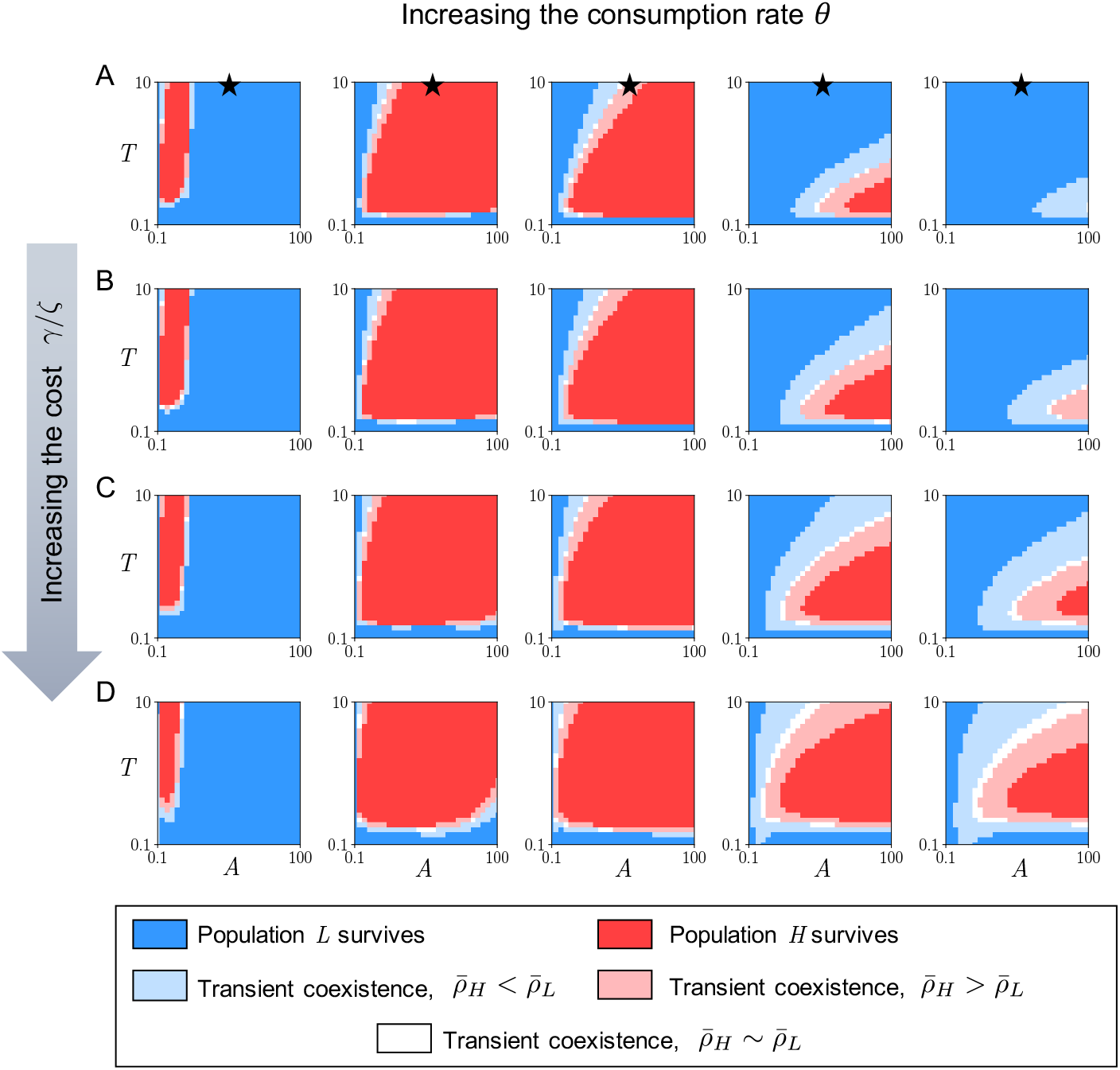
**A**. Summary of the results of numerical solutions of Eqns. (4) and (12), with the oxygen inflow *I*(*t*) defined via Eqn. (16) for different values of *A* and *T*. Different columns correspond to different values of the consumption rate of oxygen *θ*, that is, *θ* = 2×10^−5^ (first column), *θ* = 5×10^-5^ (second column), *θ* = 10^-4^ (third column), *θ* = 5×10^-4^ (fourth column) and *θ* = 10^−3^ (fifth column). The maximum cell division rate of fully glycolytic phenotypic variants is *ζ* = *γ* and the values of the other parameters are specified in Table 1. The blue points in the *A-T* plane correspond to parameter combinations for which 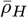 and 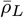 (*i.e*. the mean values of *ρ_H_*(*t*) and *ρ_L_*(*t*) computed over the last period of *I*(*t*)) are, respectively, smaller than 100 and larger than 1000 (*i.e. ρ_H_*(*t*) will eventually converge to zero), while the red points correspond to parameter combinations for which the same quantities are, respectively, larger than 1000 and smaller than 100 (*i.e. ρ_L_*(*t*) will eventually converge to zero). Moreover, the lighter regions highlight the parameter combinations for which both 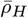 and 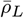, are considerably larger than 100 and 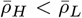 (light blue regions), 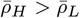 (pink regions) or 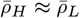 (withe regions) – *i.e*. for these parameter combinations, transient coexistence occurs for longer times although only one population will ultimately survive. The black stars highlight the parameter values corresponding to the numerical results displayed in Fig. 4. **B - D**. Same as row **A** but for *ζ* = 0.75γ (row **B**), *ζ* = 0.5γ (row **C**) and *ζ* = 0:25γ (row **D**).

The results summarised in Fig. 5 also show that, for relatively large values of *A*, when *θ* is sufficiently high (*cf*. second to fourth columns in Fig. 5), larger values of *γ/ζ* correspond to a wider range of values of the parameters *A* and *T* under which transient coexistence between the two cell populations is observed. Moreover, these results show that for sufficiently large values of *θ*, larger values of *γ/ζ* increase the likelihood that cell population *H* will ultimately outcompete cell population *L*. As illustrated by the sample dynamics presented in Fig. 6, this gives rise to smaller cell numbers, more pronounced variations in the mean phenotype of the surviving cell population and higher levels of phenotypic heterogeneity. On the other hand, the plots in the first column of Fig. 5 show that for relatively small values of *A* and *θ* the outcome of the competition between the two cell populations is only weakly affected by the quotient *γ/ζ*.

**Fig. 6.**
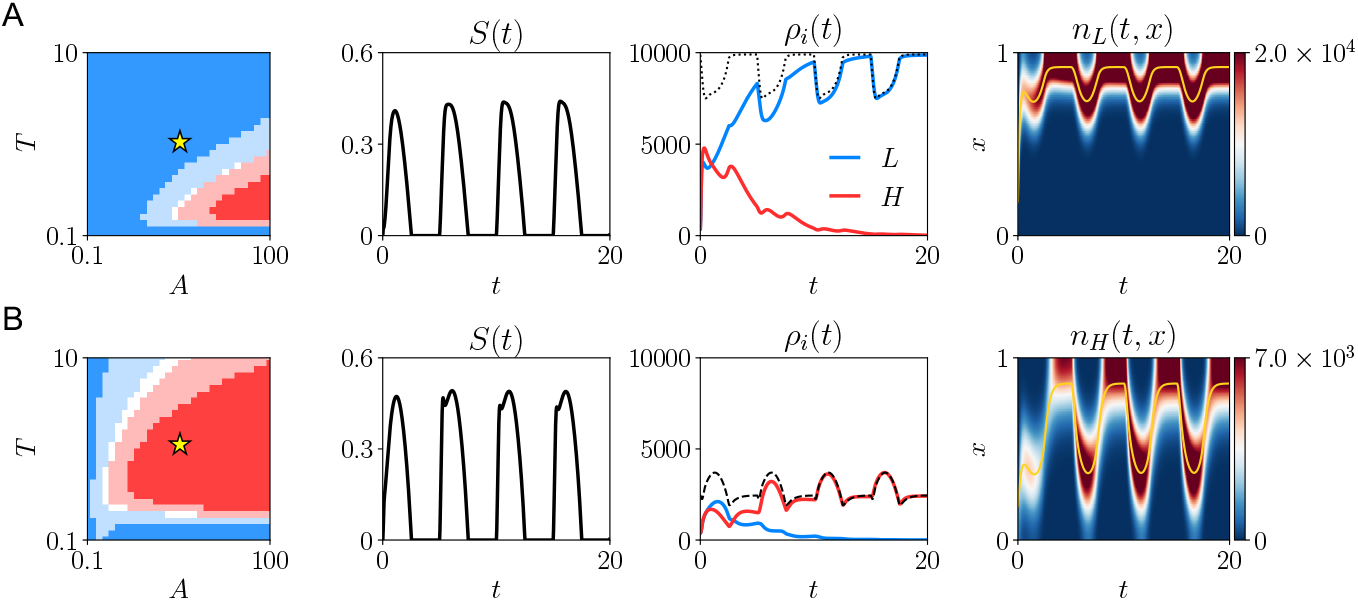
**A**. The plots in the first, column are the same as the plots in the fourth column of Fig. 5A (row **A**) and Fig. 5**D** (row **B**), and the yellow stars highlight the parameter values corresponding to the numerical results displayed here. Dynamics of the oxygen concentration *S*(*t*)(second column), the population sizes *ρ_H_*(*t*) (third column, red line) and *ρ_L_*(*t*)(third column, blue line), and the population density function of the surviving population *n_i_*(*t, x*) (fourth column) obtained by solving numerically Eqns. (4) and (12), with oxygen inflow *I*(*t*) defined via Eqn. (16) with *A* = 50 and *T* = 5. The dotted (or dashed) lines in the third column highlight the *T*-periodic solution 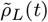 (or 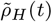) of the problem (A.11) in Appendix A, while the yellow lines in the fourth column highlight the mean phenotype *μ_i_*(*t*). The consumption rate of oxygen is *θ* = 5 × 10 the maximum cell division rate of fully glycolytic phenotypic variants is *ζ* = *γ*, and the values of the other parameters are defined as in Table 1. **B** Same as row **A** but for *ζ* = 0.25γ.

Taken together, these results indicate that, when oxygen inflow undergoes periodic oscillations, chronic hypoxia and cycles of hypoxia followed by reoxygenation can spontaneously emerge depending on the rate at which oxygen is consumed by the cells. In this biological scenario, the evolutionary fate of cell populations that undergo heritable, spontaneous phenotypic variations at different rates depends crucially upon the rate at which cells consume oxygen, *θ*, and the fitness costs associated with glycolytic metabolism, *γ/ζ*. Overall, cell populations undergoing phenotypic variations at lower rates are to be expected to be selected when the oxygen concentration remains, on average, relatively high or under chronic-hypoxic conditions. By contrast, cell populations with higher rates of phenotypic variation will outcompete other cell populations under alternating periods of hypoxia and re-oxygenation. In the latter case, the surviving cells adopt a bet-hedging strategy switching between oxidative and glycolytic metabolic phenotypes. Moreover, when oxygen levels fluctuate between zero and sufficiently larger values, higher *θ* and *γ/ζ* can favour the transient coexistence of competing populations of cells that undergo heritable, spontaneous phenotypic variations at different rates.

## 4 Application of the results to the emergence of phenotypic heterogeneity in vascularised tumours

In small tumours, cancer cells can acquire growth factors and nutrients through diffusion from blood vessels in adjacent normal tissue. However, this only supports tumour growth to a diameter of a few millimetres. Further expansion requires intratumoural blood flow and, therefore, selects for cancer cells with an ‘angiogenic’ phenotype. This results in growth of blood vessels (*i.e*. sprouting angiogenesis) into the tumour. However, unlike normal tissue, a tumour cannot perform the coordinated functions required for vascular maturation. Thus, while intratumoural blood flow does enable the transport of fresh nutrients into the tumour (Carmeliet and Jain, 2000), the tangled, uncoordinated vascular structure typically results in regions of chaotic blood flow with stochastic (but often frequent) changes in microenvironmental conditions. Thus, cancer cells at tumour-host interface, which invade adjacent normal tissue and often transiently acquire normal vessels, may have relatively stable environments, (*cf*. the left region of the scheme displayed in Fig. 7), whereas deeper regions of the tumour, characterised by limited oxygen diffusion, bring about chronic hypoxia (*cf*. the right region of the scheme displayed in Fig. 7). Moreover, regions that require angiogenesis are often subject to variable blood flow and associated microenvironmental conditions. Nonlinear interplay between vascular remodelling associated with on-going angiogenesis and oxygen consumption by the cells brings about alternating periods of hypoxia and re-oxygenation in vascularised regions in the interior of the tumour (*cf*. the central region of the scheme displayed in Fig. 7).

**Fig. 7.**
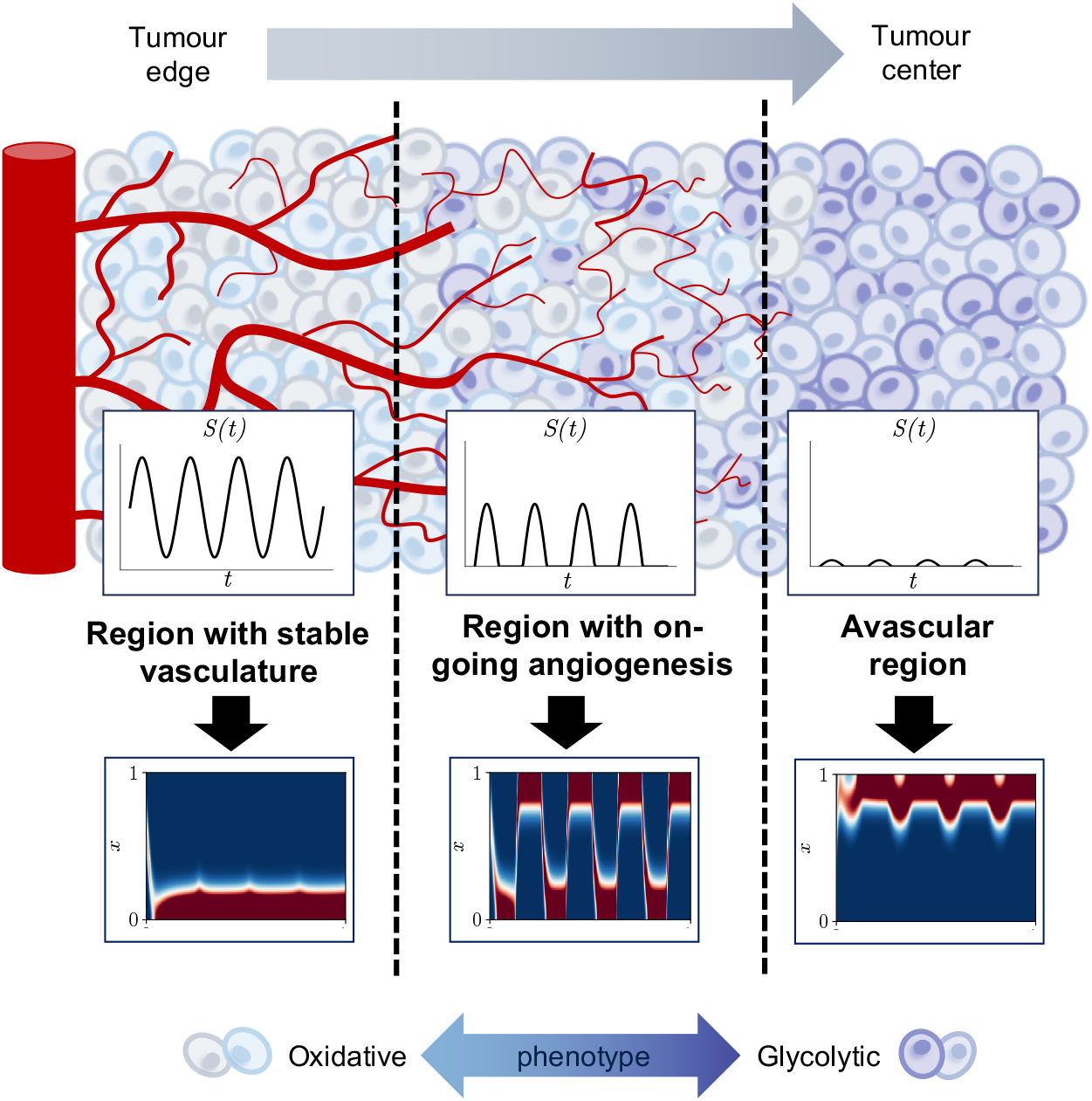
Application of the results to the emergence of phenotypic heterogeneity in vascularised tumours.

The results of our theoretical study indicate that such an expected spatiotemporal variability in oxygen concentration across tumour can create distinct ecological niches in which different phenotypic variants undergoing heritable, spontaneous phenotypic variations at different rates can be selected, and that this can also foster the emergence of phenotypic intratumour heterogeneity (*cf*. the plots of the phenotype distributions *n_i_*(*x, t*) in the lower part of Fig. 7). In particular, cell populations characterised by lower rates of phenotypic variation and a more oxidative metabolism can be expected to colonise the oxygenated regions with relatively stable vasculature at the edge of tumour; cell populations characterised by higher rates of phenotypic variation that switch between oxidative and glycolytic metabolism are likely to populate regions of on-going angiogenesis at an intermediate distance from the tumour edge; cell populations characterised by lower rates of phenotypic variation and a more glycolytic metabolism can be expected to colonise central, avascular regions of the tumour where chronic-hypoxia occurs.

## 5 Conclusions and research perspectives

Cancer cells, like all living systems, are subject to Darwinian dynamics that require them to continuously adapt to environmental conditions. Within each cancer, the micro-environmental selection forces can vary spatially due to regional variations in blood flow. However, most cancers are highly dynamic structures so that conditions within each region can also vary with time caused by variations in blood flow within a disorganised intratumoural vascular network. Prior theoretical studies have suggested temporal variations in environmental conditions may apply selection forces that result in cellular- and population-level dynamics that have significant and highly negative clinical consequences (Gravenmier et al., 2018; Gillies et al., 2018). In this work, we have developed a mathematical modelling approach to investigate the optimal adaptive strategies for cancer cells when subject to constant and periodically-oscillating oxygen inflow.

For both cases, there is excellent agreement between numerical simulations of our model and analytical results from a simplified model, which is based on asymptotic analysis of evolutionary dynamics carried out in Ardaševa et al. (2020) (see Appendix A). This agreement shows both the robustness of the biological conclusions drawn from the simulation results and the idea that the key features of the analytical results that we derived previously carry through when additional biological complexity is incorporated into the model. Furthermore, because our results persist across a range of values of the consumption rate of oxygen, *θ*, and the fitness cost associated with glycolytic metabolism, *γ/ζ*, we conclude that they are applicable to a variety of cancer cell lines under different environmental conditions, such as different levels of acidity (Casciari et al., 1992).

In summary, the simulation results generated from our model indicate that nonlinear interactions between cells and oxygen can lead naturally to the occurrence of chronic hypoxia and cycles of hypoxia and re-oxygenation depending on the rate at which oxygen is consumed by the cells. Moreover, the model supports the idea that under chronic-hypoxia lower rates of phenotypic variation constitute a source of competitive advantage. On the other hand, higher rates of phenotypic variation can confer a competitive advantage under time-varying oxygen levels, when the fitness costs associated with glycolytic metabolism are higher. In this case, the model demonstrates that bet-hedging strategies, where cells switch between oxidative and glycolytic metabolic phenotypes, can spontaneously emerge. This provides a theoretical basis for previous experimental results, such as those presented by Verduzco et al. (2015) and Chen et al. (2018), showing that intermittent hypoxia can trigger the emergence of different phenotypic properties in cancer cell populations. These results support the concept of ‘vascular normalisation’ to stabilise the cancer environment as a key strategy in cancer treatment (Jain, 2001, 2005). Furthermore, in line with previous theoretical studies indicating that periodically fluctuating environments can promote coexistence of competing populations (Hastings, 2004), our results suggest that, when the environmental conditions within the tumour switch between oxygen-poor and oxygen-rich, higher rates of oxygen consumption by cells and higher fitness costs associated with glycolytic metabolism can promote transient coexistence of competing cell populations that undergo heritable, spontaneous phenotypic variations at different rates. Finally, we have discussed how our mathematical results can shed light on the evolutionary process underlying the emergence of phenotypic heterogeneity in vascularised tumours.

We conclude with an outlook on possible extensions of the present work. The first natural extension would be to model the interplay between spontaneous and stress-induced phenotypic variations which is likely to drive cell adaptation to faster oxygen fluctuations, such as those underlying cycling hypoxia. Moreover, it would be useful to describe the metabolic dynamics of the cells in greater detail. It would also be interesting to model explicitly the evolution of the concentrations of glucose and lactic acid. In the same way, it would be interesting to extend the model to account for dynamics of reactive oxygen species that promote DNA damage and lead to mutagenesis (Liou and Storz, 2010).

Building upon previous work on the derivation of deterministic continuum models for the evolution of populations structured by phenotypic traits from stochastic individual-based models (Champagnat et al., 2002, 2006; Chisholm et al., 2016; Stace et al., 2019), it would also be interesting to develop a stochastic individual-based model corresponding to the continuum model presented here. This would make it possible to explore the impact of stochastic fluctuations in single-cell phenotypic properties on the outcome of the competition between cell populations undergoing phenotypic variations at different rates. Such stochastic effects are expected to be relevant in the regime of low cell numbers and cannot easily be captured by continuum models like the one considered here.

An additional development of our study would be to incorporate into the model spatial structure, as done for instance by Lorz et al. (2015) and Lorenzi et al. (2018), and to distribute multiple blood vessels across the spatial domain, as done for instance by Villa et al. (2019). We could then allow the formation of new blood vessels via angiogenesis, which is known be triggered by hypoxia (Dong et al., 2019). This would enable a more detailed assessment of the way in which the interplay between spatial and temporal variability of oxygen levels may dictate the phenotypic composition and the level of phenotypic heterogeneity of vascularised tumours. Moreover, since experimental results suggest that cycling hypoxia increases cell motility and promotes the formation of metastases (Liu et al., 2017; Chen et al., 2018), when including spatial structure in the model it would also be interesting to explore the adaptive role of the trade-off between cell motility and cellular proliferation (Gallaher et al., 2019). Such a model would have the potential to inform new treatment strategies aimed at minimising the pro-metastatic effect of cycling hypoxia.

## A Analysis of evolutionary dynamics for a simplified model

In order to obtain a detailed analytical description of the evolutionary dynamics of the two cell populations, we can consider a simplified scenario whereby the ODE for *S*(*t*) is decoupled from the system of non-local parabolic PDEs (4). In particular, we let the evolution of the oxygen concentration *S*(*t*) be governed by the following Cauchy problem

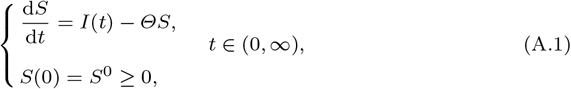

where the effects of oxygen consumption and oxygen decay are both encapsulated in the parameter *Θ* > 0. Moreover, to facilitate analysis, we extend the interval [0, 1] to 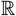 and re-define the population-level quantities accordingly, *i.e*. we use the definitions

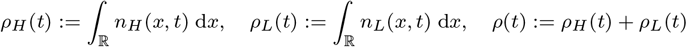

and

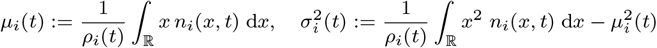

with *i* ∈ {*H, L*}. Finally, in agreement with much of the previous work on the mathematical analysis of the evolutionary dynamics of continuous traits, which relies on the simplifying assumption that population densities are Gaussians (Rice, 2004), we consider initial conditions of the form

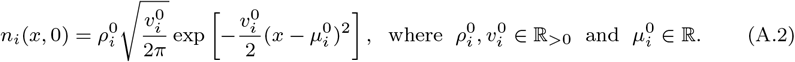

This allows us to use the result established by Proposition 1, which can be proved through the method that we previously employed in Ardaševa et al. (2020).

### Proposition 1

*Under assumptions* (6) *and* (7), *the system of non-local PDEs* (4) *posed on* 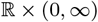 *and subject to the initial condition* (A.2) *admits the exact solution*

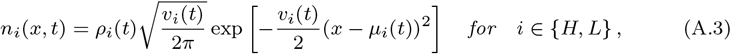

*with the population size, ρ_i_*(*t*), *the mean phenotype, μ_i_*(*t*), *and the inverse of the phenotypic variance*, 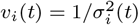, *being solutions of the Cauchy problem*

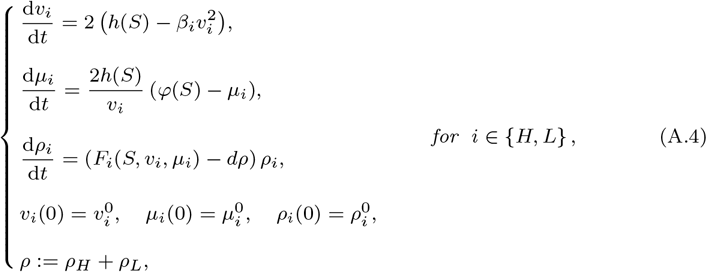

*where*

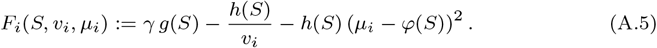

In the case where the inflow of oxygen is constant, *i.e*. the source term *I*(*t*) in the ODE (A.1) satisfies assumption (14), our main results are summarised by Theorem 1, where the functions *g, φ* and *h* are defined according to (9) and (10), and we use the definitions

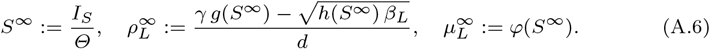

### Theorem 1

*Under assumptions* (5)-(11) *and the additional assumption* (14), *the solution of the system of non-local PDEs* (4) *posed on* 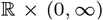, *subject to the initial condition* (A.2) *and complemented with the Cauchy problem* (A.1) *is of the Gaussian form* (A.3) *and satisfies the following:*

i. *if*

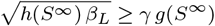

*then*

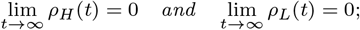
ii. *if*

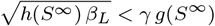

*then*

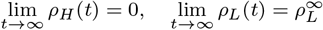

*and*

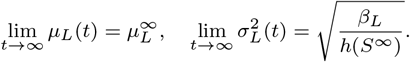

In the case where the inflow of oxygen undergoes periodic oscillations, *i.e*. the source term *I*(*t*) in the ODE (A.1) satisfies assumption (15) along with the additional assumption

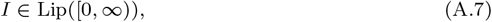

our main results are summarised by Theorem 2, where 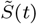 is the unique non-negative *T*-periodic solution of the problem

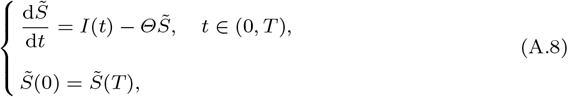

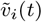 is the unique real *T*-periodic solution of the problem

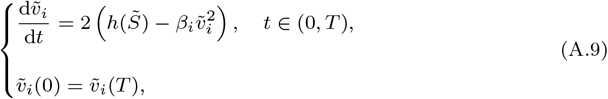

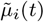 is the unique real *T*-periodic solution of the problem

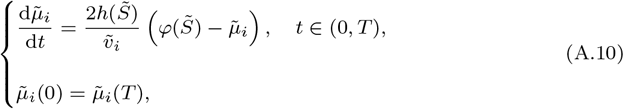

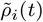 is the unique real non-negative *T*-periodic solution of the problem

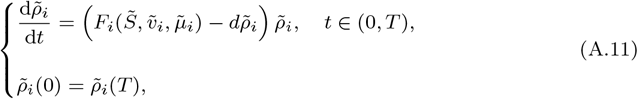

and

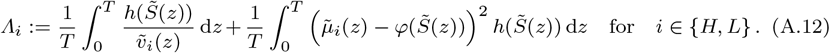

In (A.9)-(A.12), the functions *g, φ* and *h* are defined according to (9) and (10). Moreover, the function *F_i_* in (A.11) is defined according to (A.5).

### Theorem 2

*Under assumptions* (5)-(11) *and the additional assumptions* (15) *and* (A.7), *the solution of the system of non-local PDEs* (4) *posed on* 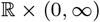, *subject to the initial condition* (A.2) *and complemented with the Cauchy problem* (A.1) *is of the Gaussian form* (A.3) *and satisfies the following:*

i. *if*

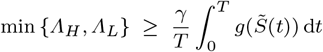

*then*

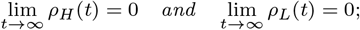
ii. *if*

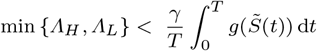

*and*

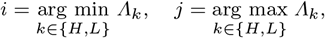

*then*

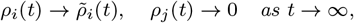

*and*

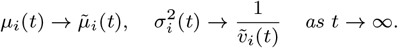

Theorem 1 and Theorem 2 can be proved through methods similar to those that we employed in Ardaševa et al. (2020) and, therefore, their proofs are omitted here.

